# Proteomics to uncover new actors in the antifungal action of metformin. Impact on virulence traits, oxidative stress, and essential proteins from *Candida albicans*

**DOI:** 10.64898/2026.09.11.750873

**Authors:** Victoria Mascaraque, Celia Ramos, Cristina Navas, María Luisa Hernáez, Concha Gil, Gloria Molero

## Abstract

Metformin is one of the most widely prescribed drugs with a proven safety profile, making it an excellent candidate for drug repurposing strategies. Its anti-*Candida* effect and its synergistic potential with azoles have already been described against *C. albicans* and *C. glabrata*, alongside AMPK involvement and its divergent consequences on autophagy. However, metformin has additional effects on *C. albicans* biology that remain to be explored using differential proteomics. In the present study, *C. albicans* SC5314 was treated with metformin to identify the proteins involved in its anti-*Candida* activity. First, we demonstrated that metformin inhibits *C. albicans* SC5314 growth in a dose-dependent manner, an effect that intensifies under low glucose (0.2%) and filament-inducing conditions. It also reduces virulence traits such as filamentation, biofilm formation, and invasive growth. Furthermore, it induces significant oxidative stress, which is neutralized by antioxidant agents such as N-acetylcysteine and glutathione. Additionally, 50 mM metformin substantially increased fluconazole efficacy under 0.2% glucose. Our label-free proteomic study of *C. albicans* SC5314 exposed to 50 mM metformin allowed the identification and quantification of 1,899 proteins, with 95 and 47 proteins showing increased and decreased abundance, respectively. Notably, 26 of the down-regulated proteins were encoded by essential genes, demonstrating the drastic effect of metformin on *C. albicans* viability. GO Term analysis revealed that the most relevant functions affected were ATP binding, inhibition of ATPase activity (with reduced ATP levels), translation inhibition, mitochondrial function, filamentation, and responses to oxidative stress and antifungals. In conclusion, metformin significantly compromises the viability and virulence of *C. albicans* at high concentrations. Given that high concentrations of metformin have recently been reported in the intestinal tract, and that intestinal *Candida* is a known source of invasive infections in immunocompromised, under chemotherapy, and post-surgery patients, understanding metformin’s anti-*Candida* action becomes increasingly relevant.

## Introduction

*Candida albicans* is an opportunistic pathogen and one of the most important fungal pathogens that compromise human health, causing superficial infections and life-threatening invasive candidiasis in immunocompromised patients [1]. Invasive candidiasis remains the most common cause of invasive fungal infections worldwide, with more than 1.5 million cases and almost 1 million deaths per year. As fungal infections are a growing concern worldwide, the World Health Organization elaborated the Fungal Priority Pathogen List, including *C. albicans* in the critical group, owing to the annual increase in invasive candidiasis incidence [2, 3]. The source of many *Candida* infections are colonized skin, mucosa, or the gastrointestinal tract. Damages to the GI mucosa by chemotherapy (mucositis), trauma, or sepsis allow the penetration of microorganisms into the injured tissue and gain access to the lymphatics and bloodstream [4-6].

A relevant problem in candidiasis is the limitation of antifungal treatments against *Candida* infections, together with an increase in resistance to antifungal agents [7]. Thus, the discovery of new antifungal targets and agents remains an important task, in addition to drug repurposing, which is proving to be a promising solution [8, 9]. In this context, the antihyperglycemic drug metformin has emerged as a potential antifungal agent against *Candida* species [10, 11], and we have explored its mechanism of action and potentiation of other antifungal activities [10].

Metformin is one of the most widely prescribed drugs worldwide, mainly for the treatment of type II diabetes [12]. When administered orally, the usual dose of metformin (0.5–3= g/day) is effective for long-term glycemic control, with notable changes in HbA1c [13, 14]. Potential new indications for metformin have been proposed, such as the inhibition of tumor cell proliferation, cardiovascular and neurological protection, treatment of polycystic ovary syndrome [15], and anti-inflammatory properties that help reduce mortality rates due to COVID-19 infection and the development of prolonged symptoms [16].

However, the mechanisms underlying the therapeutic action of metformin are complex and still not fully understood. Furthermore, they seem to vary depending on the dose, duration of treatment [17], and niche [18, 19]. The therapeutic plasma concentrations are approximately 10-70 µM, and the accepted mechanism of action is the activation by phosphorylation of AMPK (AMP-activated protein kinase)[20]. Subsequently, AMPK-independent mechanisms have also been observed [21]. Mitochondria have been described as the main organelle target, specifically by the inhibition of the mitochondrial respiratory chain complex I [22], although inhibition of complex IV has also been suggested [23]. In addition, the inhibition of v-ATPase in lysosomes by low concentrations of metformin has been proposed as a new mechanism of action [24]. The metformin effect seems to be pleiotropic, as a recent study on JHH-7 cells showed effects on protein turnover, ubiquitination, DNA damage, and the cell cycle [25]. The authors also used proteomics to demonstrate these effects. With respect to the concentrations used, Sebo et al. (2026) have shown that the concentration of metformin is 30-300 times higher in the intestine than in plasma, where it inhibits the mitochondrial respiratory chain complex I from the intestinal epithelial cells [26].

Regarding its antimicrobial activity, metformin has been used as an antiparasitic agent with promising results [27, 28]. In addition, its antibacterial and antiviral properties have been reported [29-32]. With respect to antifungal activity, Xu et al. (2018) described the effects of several biguanides, including metformin, on *Candida* species, focusing on *C. glabrata*. They also demonstrated the enhancement of azole and amphotericin B activity in combination with metformin against *C. glabrata*. Recently, Zhao et al. (2025) compared the effects of metformin on the growth of *C. albicans* in planktonic and biofilm states and observed an antifungal mechanism through autophagy modulation.

In the present study, we analyzed the phenotypic effects of metformin on the *C. albicans* wild-type strain SC5314 and the proteomic changes induced by label-free quantitative proteomics, with a focus on discovering new antifungal mechanisms. Metformin has significant effects on *C. albicans* biology and induces important changes in proteins related to translation, dimorphic transition, antifungal response, oxidative stress response, and ATP synthesis coupled with electron transport. In addition, more than half of the proteins with decreased abundance after treatment were encoded by essential genes.

## Experimental Procedures

### Experimental Design and Statistical Rationale

In this study, we have analyzed the phenotypic effects of increasing concentrations of metformin by testing different culture media, temperatures, and glucose concentrations. Once the best conditions were established, a label-free proteomic approach was used to analyze the proteins that varied in their quantities in response to metformin compared to the untreated control. Functional in silico studies were also performed. Based on these results, mutants in some of the most interesting pathways affected were analyzed, together with measurements with reagents affecting important cellular functions, to show possible new mechanisms of action of metformin.

All experiments were performed with independent biological replicates, with specific n values provided in the corresponding figure legends. Statistical analyses were conducted using two-tailed Student’s t-tests or one-way ANOVA, where appropriate, with p < 0.05 considered statistically significant. Data are presented as mean ± standard deviation (SD) unless otherwise specified. Further details on the experimental procedures and data analysis pipelines are provided in the following sections.

### Strain, media and growth conditions

*C. albicans* SC5314 [33] was used as the wild type, and several mutants from Noble’s collection [34] were used for the specific tests. Yeast cells were pre-cultured in liquid YPD medium (10 g/L yeast extract, 20 g/L peptone, and 20 g/L glucose) in a rotary shaker at 180 rpm and 30°C overnight, unless otherwise indicated. When used, RPMI 1640 with 2 mM L-glutamine and 2 g/L glucose (Lonza) was supplemented with antibiotics (penicillin 10000 U/ml, streptomycin 10000 U/ml) and 10% heat-inactivated fetal bovine serum (FBS). Fluconazole was obtained from Sigma-Aldrich and dissolved in dimethyl sulfoxide (DMSO). Metformin hydrochloride (1,1-Dimethylbiguanide hydrochloride) was purchased from ACROS Organics and dissolved in ultrapure water to prepare a 500 mM (85.40 mg/mL) stock solution immediately before use.

### Metformin susceptibility testing

Metformin susceptibility was assessed using microdilution plate assays. Briefly, yeast cells pre-cultured in liquid yeast extract-peptone-dextrose (YPD) medium were washed twice with ice-cold phosphate-buffered saline (PBS) and resuspended in the medium used for the assay (complete RPMI or YPD). The final inoculum was 2.5 × 10^5^ CFU/mL, and the highest concentration tested was 100 mM metformin (17.08 mg/mL). The total volume per well was 150 µL. The cells were incubated at 37°C for 20-24 h. After incubation, turbidity was observed, and OD_595nm_ was measured using a microdilution plate reader. Cell morphology was observed using a Nikon Eclipse TE2000-U microscope connected to a high-resolution Hamamatsu ORCA-ER camera system. To test viability, 5 µL of 10-fold dilutions from the first four wells with the highest concentrations of metformin were spotted on YPD agar plates and incubated for 24h at 37°C. When necessary, hydrogen peroxide, menadione, sorbitol, KCl (Sigma-Aldrich), and NaCl (PanReac AppliChem) were added to agar plates.

When growth curves were obtained, round-bottom polystyrene 96-well microtiter plates were used, with a total volume of 180 µL per well and 1.7 × 10^6^ CFU/mL as the final inoculum. Cells were incubated at 37°C for 24 h with shaking in a SPECTROstar^Nano^ (BMG LABTECH), and the optical density at 600 nm was monitored every 30 min. When necessary, puromycin, N-acetylcysteine, or glutathione (Sigma-Aldrich) were added.

### *In vitro* adhesion capacity

*Candida* cells (2.5 × 10^5^ CFU/mL) were incubated in complete RPMI medium with different concentrations of metformin in flat-bottom polystyrene 96-well microtiter plates overnight. The cells were then stained with 25 µL of crystal violet solution (0.1% in ethanol) per well for 5 min. Finally, non-adherent cells and excess crystal violet were removed by washing with water, and the stained cells were observed using microscopy, as mentioned before.

### Agar invasion assays

For invasive growth on solid media, cells were plated onto Spider agar (10 g/L nutrient broth, 10 g/L mannitol, 2 g/L K2HPO4, 11 g/L agar) supplemented with different concentrations of metformin at a density of 30-40 CFU/plate. The plates were incubated at 37°C for 7 d, and fungal colonies were photographed using a Wild Heerbrugg M5-46860 Stereo Microscope coupled to a digital camera (Panasomic Lumix DMC-G1K).

### Cultures and cell extracts for Proteomics Analysis

*C. albicans* SC5314 (2.5 × 10^6^ CFU/mL) control and treated with 50 mM metformin were grown in 50 mL complete RPMI culture at 60 rpm and 37°C for 6 h. Cells were harvested by centrifugation at 2500 rpm for 5 min at 4°C in an Eppendorf centrifuge 5810R, washed with PBS, and frozen at -80°C until use. Four biological replicates were performed under each condition. Cell extracts were obtained by suspending cells in 200 µL of cold lysis buffer (50 mM Tris-HCl pH 7.5, 1 mM EDTA, 150 mM NaCl, 1 mM DTT, 0.5 mM PMSF, and 1% protease inhibitor cocktail (PierceTM)). Subsequently, an equal volume of 0.5-0.75 mm diameter glass beads was added, and cells were disrupted mechanically in a FastPrep-24TM (MP Biomedicals) by applying eight 20s rounds at 5.5 speed with intermediate ice cooling. Cell extracts were clarified by centrifugation at 13000 rpm for 15 min at 4°C, and the supernatants were collected and stored at -80°C. Protein concentration was measured using the Bradford protein assay, and the protein pattern of the samples was compared using SDS-PAGE in a 10% polyacrylamide gel stained with Coomassie Brilliant Blue G250.

### Quantitative Proteomic Analysis

#### Sample preparation and digestion

Proteomic analysis was performed at the Proteomics Unit of Complutense University of Madrid. Proteins from cell extracts were precipitated using the methanol/chloroform method [35] and resuspended in 100-150 µL of 8 M Urea. Each sample (50 µg) was reduced with 10 mM DTT at 56°C for 30 min and alkylated with 55 mM iodoacetamide for 20 min in the dark. Subsequently, the proteins were digested with 1 µg recombinant trypsin (Roche Molecular Biochemicals) in 25 mM ammonium bicarbonate pH 8.5 at 37°C overnight. After digestion, the peptides were acidified, desalted, and concentrated by reverse-phase chromatography C18 onto Bond Elut OMIX tips (Agilent Technologies). The peptides were then dried in a Savant SpeedVac (Thermo Fisher Scientific), resuspended in 12 µL of 2% acetonitrile and 0.1% formic acid, and quantified using a Qubit 3.0 fluorometer (Thermo Fisher Scientific).

#### LC-MSMS analysis

Peptides (1.2 µg) from each sample were analyzed by nano-liquid chromatography using an Easy-nLC 1000 System (Thermo Scientific) coupled to a Q-Exactive HF mass spectrometer (Thermo Scientific). Peptides were loaded onto an Acclaim PepMap 100 Trapping column (Thermo Scientific, 20 mm × 75 µm ID, 3 µm particle size C18 resin with 100 Å pore size) and then separated and eluted on a C18 resin analytical column Picofrit (Thermo Scientific Easy Spray Column, PepMap RSLC C18, 500 mm × 75 µm ID, 2 µm particle size resin with 100 Å pore size) with an integrated-spray tip. A 180 min gradient of 2% to 40% buffer B (0.1% formic acid in acetonitrile) in buffer A (0.1% formic acid in water) at a constant flow rate of 250 nL/min was employed. Data acquisition was performed using a Q-Exactive HF mass spectrometer. Data were acquired using an ion spray voltage of 1.8 kV and an ion transfer temperature of 270°C. All data were acquired using data-dependent acquisition (DDA) in the positive ion mode. Full-scan MS spectra (m/z 350-2000 Da) were acquired at a resolution of 60000, and the 15 most intense ions with charges of 2 to 6+ were selected for High Collision Dissociation (HCD) fragmentation with a normalized collision energy of 27% and a dynamic exclusion of 27s.

#### Proteins identification and label-free quantification

Peptide identification from raw data was performed using the MASCOT v.2.6.1 search engine through Proteome Discoverer 2.2 software (Thermo Fisher Scientific). A database search was performed against CGD21 (6279 sequences). The search parameters included tolerances of 10 ppm for precursor ions and 0.02 Da for MS/MS fragment ions, two missed cleavages allowed, carbamidomethylation of cysteines as a fixed modification, and oxidation of methionine as a variable modification. The identified peptides were validated using the Percolator algorithm with a q-value threshold of 0.01. A nested design was used to compare the samples from cells treated with metformin and the control cells. Peptide label-free quantification was performed using Proteome Discoverer v2.2 with node precursor ion quantification. The peptide ratios were normalized to the geometric median of all ratios in each biological replicate. The protein ratios were determined as the geometric median of all quantified peptides belonging to a specific protein. Further analysis was performed only with proteins with the following additional acceptance criteria: proteins identified in at least four of the eight samples when detected in both conditions (control and metformin treatment) and in at least three samples when detected only in one condition; proteins with at least two peptides quantified without discrepant values inside each protein; proteins with only one peptide quantified when peptide-spectrum match (PSM) was more than two and without discrepant values inside each peptide; and proteins with a q-value less than 0.05.

All mass spectrometry proteomics data have been deposited in the PRIDE repository [36] with the dataset identifier PXD014456.

#### GO term enrichment and bioinformatic analysis

Gene Ontology (GO) enrichment analysis was performed using the GO Term Finder tool from *the Candida* Genome Database [37]. Volcano plot and Venn diagram were generated using SRplot [38]. STRING software version 12.0 (https://string-db.org) was used to study protein-protein interactions [39].

### Measurement of the intracellular ATP concentration

Intracellular ATP levels in C. albicans cultures were measured colorimetrically using an ATP Assay Kit (Sigma-Aldrich). Briefly, *C. albicans* SC5314 (2.5 × 10^6^ CFU/mL) control and treated with 50 mM metformin were grown in 10 mL complete RPMI cultures at 60 rpm and 37°C for 3 h. Cells were harvested by centrifugation at 2500 rpm for 5 min at 4°C in an Eppendorf centrifuge 5810R and washed thrice with PBS. Cells were disrupted in 250 µL of ATP Assay Buffer using a High Intensity Ultrasonic Processor VC375 (Sonics & Materials) for 20 cycles of 10 s, with cooling between cycles, at a 40% duty cycle and 4 output controls. Cell lysates were centrifuged to remove the remaining cell debris and then deproteinized using a 10 kDa MWCO spin filter (Millipore). The assay was performed following the manufacturer’s instructions, incubating the microdilution plate with gentle shaking (300 rpm) in a ThermoMixer (Eppendorf) at 22°C (room temperature) for 30 min in the dark. Absorbance was measured at 570 nm using an SPECTROstar^Nano^ (BMG LABTECH), and ATP concentrations were determined from ATP standard curves.

## Results

### Metformin affects *C. albicans* viability, morphogenesis and biofilm formation

The effect of metformin on *C. albicans* SC5314 growth strongly depended on the concentration and the culture medium used. As shown in **Fig. 1**, growth was inhibited only at 50 and 100 mM concentrations in complete RPMI culture medium **(Fig. 1A-B)** and with less intensity in the traditional YPD medium containing glucose 2% **(Fig. 1C-D)**, in both cases, at 37°C. The effect of metformin was lower when the stock solution was not prepared immediately before use; therefore, metformin was freshly prepared for all experiments. It must be underscored that *C. albicans* was able to filament in complete RPMI culture medium but not in YPD under the assayed conditions. Some virulence traits, such as filamentation, invasive growth, and biofilm formation, were analyzed. As can be observed in Fig. 1, the filamentation capacity was impaired in the presence of 50 mM metformin, and the filaments have a different morphology **(Fig. 1E)**. In Spider medium, *C. albicans* was able to grow even in the presence of 100 mM metformin; however, the colony size and invasive growth were impaired **(Fig. 1F)**. With respect to biofilm formation, crystal violet staining showed a reduction in the biofilm matrix, even in the presence of low concentrations of metformin **(Fig. 1G)**.

**Fig. 1.**
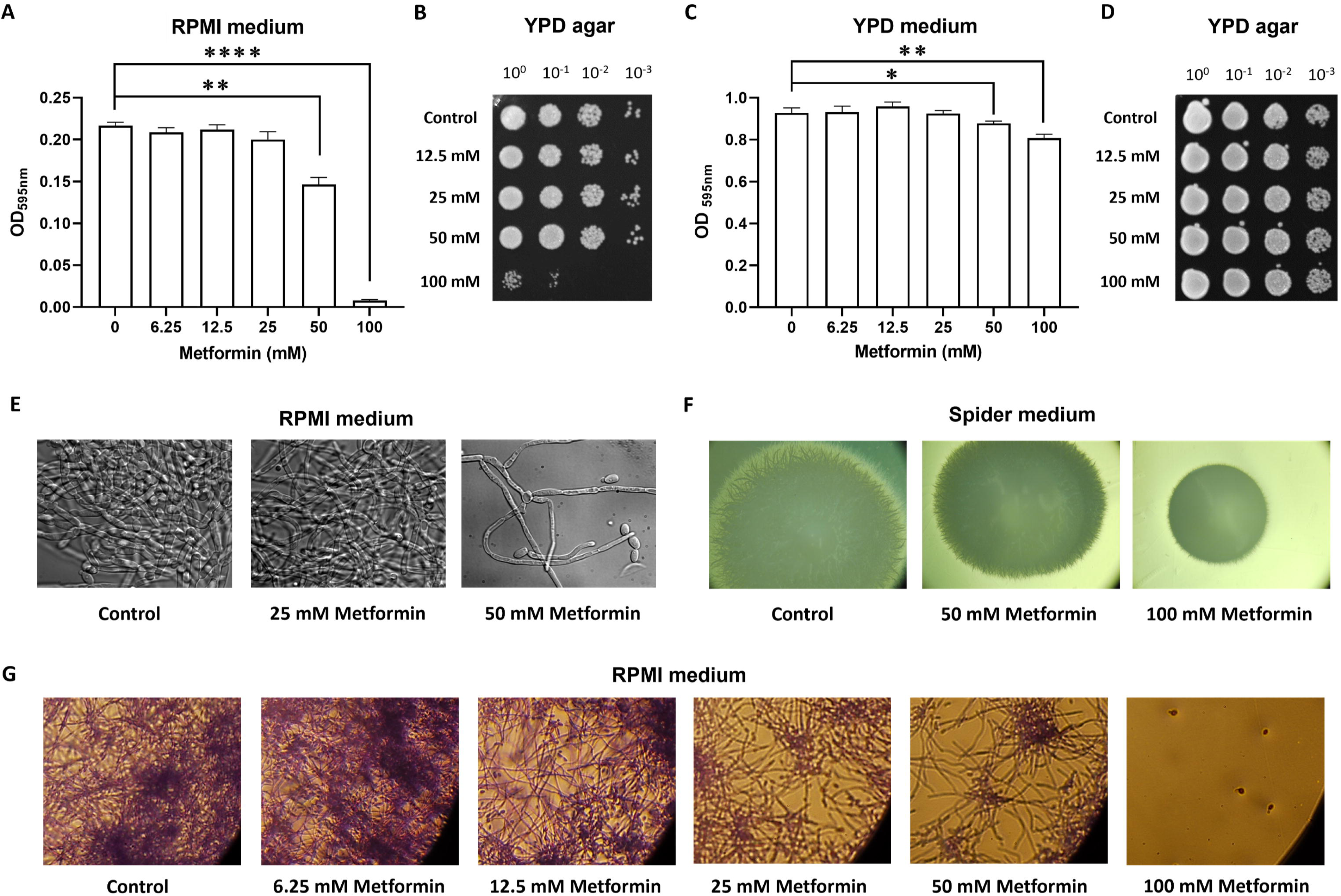
Phenotypic effects of metformin. **(A)** Sensitivity to increasing concentrations of metformin in complete RPMI in microdilution plates and **(B)** cell viability of ten-fold serial dilutions on YPD agar plates after treatment in RPMI. **(C)** Sensitivity to increasing concentrations of metformin in YPD medium in microdilution plates and **(D)** cell viability of ten-fold serial dilutions on YPD agar plates after treatment; \**p* < 0.05, \*\**p* < 0.01, \*\*\*\**p* < 0.0001, unpaired t-test. **(E)** Cell morphology observed by microscopy after treatment in microdilution plates with increasing concentrations of metformin in complete RPMI. **(F)** Agar invasion assay in Spider medium supplemented with different concentrations of metformin. Fungal colonies were photographed using a 25X objective lens. **(G)** Crystal violet staining after treatment in microdilution plates with increasing concentrations of metformin in complete RPMI. All experiments were performed at an incubation temperature of 37 °C.

As glucose concentration is important for metformin action [40, 41], growth curves of *C. albicans* were obtained in complete RPMI and YPD with 0.2% or 2% glucose **(Supplementary Fig. S1)**, showing that when glucose is increased in RPMI medium, the 50 mM metformin effect on growth was lost. Conversely, in 0.2% glucose YPD, 100 mM metformin caused a 20% inhibitory effect. Therefore, the effect of metformin on *C. albicans* growth is increased under low (0.2%) concentrations of glucose.

### Proteomic changes in response to metformin

Owing to the culture medium dependence of the metformin effect, RPMI 0.2% glucose complete medium was chosen for proteomic studies. A sublethal concentration of 50 mM at 37°C for 6 h allowed us to obtain sufficient proteins for proteomic analyses. To avoid cell lysis, agitation was limited to 60 rpm. Four biological replicates of treated and control samples were used for label-free quantitative proteomics. The proteomic study allowed the identification and quantification of 1899 proteins, 142 of which had differences in abundance (q-value < 0.05), increasing 95 and decreasing 47 after the treatment **(Fig. 2 and Supplementary Tables S1 and S2)**.

**Fig. 2.**
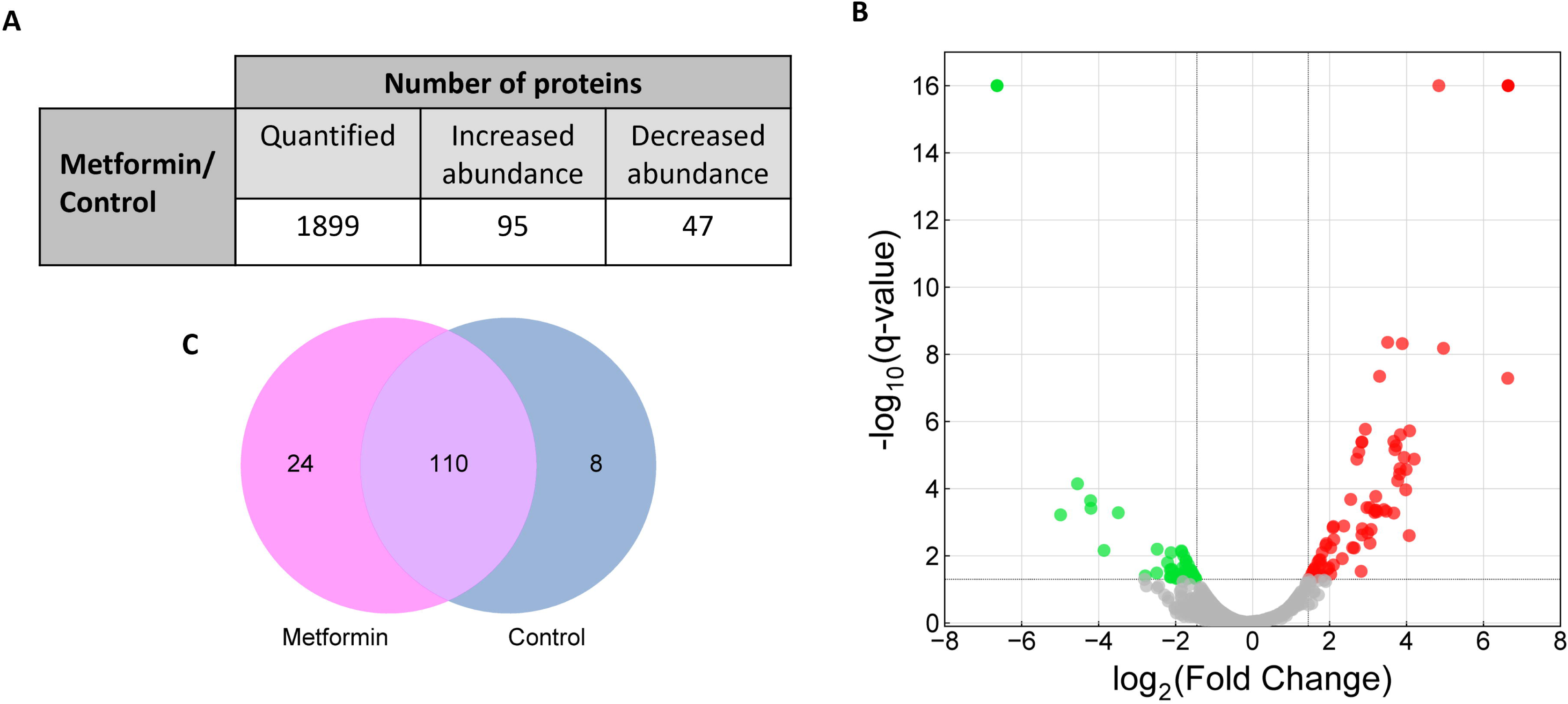
Label-free quantitative proteomics in response to metformin. **(A)** Number of quantified proteins and proteins with changes in their abundance after treatment with 50 mM metformin. **(B)** Volcano plot of quantified proteins. Changes in the abundance of proteins with a q-value < 0.05 (-log_10_ q-value > 1.3) are represented in red or green according to their increased or decreased abundance, respectively. **(C)** Venn diagram showing the number of proteins with changes in abundance.

The analysis of the proteins with the higher increases in the relative abundance after the treatment **(TABLE 1)** revealed proteins involved in ATP binding and inhibition of the ATPase activity (orf19.7345, orf19.5201.1), protein catabolism (Prb1, orf19.6753), RNases (Rnh1), ribosomal proteins (orf19.6415.1, orf19.7107, Rps30), proteins related to regulation of mRNA stability (Dcp2), DNA repair (Snf6 and Hat2), amino acid hydrolysis (Sno1), cell wall biosynthesis (Kre9), and oxidoreductase activity (Ald6 and Pst2).

**TABLE 1.** Ranking of the 15 proteins with the highest increases in relative abundance after treatment.

| Protein | Description | Metformin/Control<br>Ratio $\log_2$ |
| --- | --- | --- |
| orf19.5201.1 | Domain(s) with predicted ATPase inhibitor activity | 6.63 |
| Prb1 | Putative vacuolar protease | 4.96 |
| orf19.6415.1 | Ortholog(s) have structural constituent of ribosome activity | 4.84 |
| Rnh1 | Ribonuclease H | 4.20 |
| Dcp2 | Ortholog(s) have chromatin binding, hydrolase activity, mRNA binding activity | 4.08 |
| Snf6 | Subunit of SWI/SNF chromatin remodeling complex | 4.07 |
| Hat2 | Putative Hat1-Hat2 histone acetyltransferase complex subunit | 3.99 |
| Sno1 | Protein with a predicted role in pyridoxine metabolism | 3.98 |
| orf19.7107 | Ortholog(s) have role in ribosomal large subunit biogenesis | 3.94 |
| Rps30 | Putative ribosomal 40S subunit protein S30 | 3.89 |
| Kre9 | Protein of beta-1,6-glucan biosynthesis | 3.84 |
| orf19.6753 | Protein with a predicted RING-type zinc finger | 3.83 |
| orf19.7345 | Ortholog(s) have ATP binding activity | 3.82 |
| Ald6 | Putative aldehyde dehydrogenase | 3.77 |
| Pst2 | Putative NADH: quinone oxidoreductase | 3.73 |

Additionally, some proteins were only detected after treatment **(TABLE 2 and Supplementary Table S3)**. In these cases, the ratio between the number of peptide-spectrum matches (PSM) and the molecular weight of the protein (PSM/MW ratio) was used to rank proteins according to their relative abundance (Supplementary Table S3), indicating that the higher the protein level, the closer the punctuation to 1. This is the case for orf19.2276, whose ortholog *RPB9* in *S. cerevisiae* encodes a subunit of the RNA polymerase II core complex, and Ess1 which regulates RNA polymerase II function [42]. Furthermore, proteins related to RNA polymerase I (orf19.962), 3’-5’ RNA processing (orf19.6259), and RNA splicing (Smd2, orf19.7375) were also detected, indicating an increase in proteins related to transcription after treatment with metformin. Other interesting proteins are involved in nucleic acid biosynthesis (Met14, orf19.1110, orf19.1249, orf19.891), nucleocytoplasmic transport (orf19.6498), ribosome assembly (orf19.1642), cell wall (Rbt1), protein folding (Mdj1), post-translational modification (Bet2, orf19.6822, Uba2), as well as redox reactions (Grx3, orf19.5342) and ATP synthase complex assembly (orf19.3686).

**TABLE 2.** Proteins were identified only in metformin-treated cells.

|  | Proteins | Metformin/Control<br>Ratio log<sub>2</sub> |
| --- | --- | --- |
| Encoded by essential genes | Grx3, Ess1, Smd2, Bet2, orf19.6259, Mdj1, orf19.1110 | 6.64 |
| Encoded by nonessential genes | orf19.2276, orf19.319, orf19.4886, orf19.962, Met14, orf19.7375, orf19.1642, orf19.7443, Rbt1, orf19.6822, orf19.1249, orf19.3686, orf19.6498, Ndt80, orf19.5342, orf19.891, Uba2 |  |

Regarding the proteins with higher decreases in relative abundance after the treatment **(TABLE 3)**, the analysis revealed proteins involved in lipid organization and biosynthesis (Ist2, Asm3), nucleocytoplasmic transport (Tom1, orf19.3681), amino acid biosynthesis and transport (Thr1, orf19.5321, Gap4), protein synthesis (Wrs1, orf19.3798, Sui3), chaperones (Cct7, orf19.2720), and proteins with oxidoreductase activity (Gpx2, orf19.1480). Proteins detected only under control conditions are listed in **TABLE 4 and Supplementary Table S4**. According to the PSM/MW ratio, the protein with the highest relative decrease with respect to the treatment was related to ATPase activity (Ena21), in concordance with the inhibition of ATPase activity observed with metformin, as mentioned before. The other identified proteins participate in ER-Golgi trafficking and elimination of misfolded proteins (Erv29), oxidoreductase activity (orf19.4758), lipid biosynthesis (Slc1), protein acetylation (Naa25), ribosome biogenesis (Utp22), and rRNAs synthesis (orf19.1578).

**TABLE 3.** Ranking of the 15 proteins with the highest decrease in relative abundance after treatment.

| Protein | Description | Metformin/Control Ratio log<sub>2</sub> |
| --- | --- | --- |
| Ist2 | Ortholog(s) have lipid binding activity and role in endoplasmic reticulum membrane organization | -4.99 |
| Asm3 | Putative secreted acid sphingomyelin phosphodiesterase | -4.55 |
| orf19.5648 | Unknown protein | -4.21 |
| Tom1 | Putative E3 ubiquitin ligase | -4.20 |
| Thr1 | Putative homoserine kinase | -3.86 |
| orf19.5321 | Ortholog(s) have methylenetetrahydrofolate reductase (NAD(P)H) activity and role in methionine biosynthetic process | -3.49 |
| Gap4 | High-affinity S-adenosylmethionine permease | -2.79 |
| Wrs1* | Putative tryptophanyl-tRNA synthetase | -2.49 |
| orf19.3681* | Ortholog(s) have guanyl-nucleotide exchange factor activity | -2.48 |
| Cct7* | Cytosolic chaperonin Cct ring complex | -2.21 |
| Gpx2 | Similar to glutathione peroxidase | -2.14 |
| orf19.3798 | Ortholog(s) have tRNA (guanine(46)-N7)-methyltransferase activity | -2.14 |
| orf19.1480* | Putative succinate dehydrogenase | -2.12 |
| Sui3* | Putative translation initiation factor | -2.12 |
| orf19.2720* | Cytosolic chaperonin Cct ring complex subunit | -2.11 |

\*Proteins encoded by essential genes.

**TABLE 4.** Proteins not detected in under metformin treatment.

|  | Proteins | Metformin/Control<br>Ratio log<sub>2</sub> |
| --- | --- | --- |
| Encoded by essential<br>genes | Slc1, Utp22, orf19.1578 | -6.64 |
| Encoded by<br>nonessential genes | Ena21, Erv29, orf19.4758, Naa25, orf19.5799 |  |

Remarkably, 55% of the proteins with decreased abundance after treatment were encoded by essential genes [37, 43], showing the drastic effect of metformin on the biology of *C. albicans* **(TABLE 4 and Supplementary Table S5)**.

Gene Ontology (GO) enrichment analysis was performed to characterize proteins whose abundance significantly changed **(Supplementary Table S6)**. This study revealed that a high number of proteins related to translation, dimorphic transition, and antifungal response significantly changed their abundance in the presence of metformin.

Proteins were clustered into networks using STRING software. Proteins with significantly increased abundance and detected only after metformin treatment were grouped into nine clusters, while those with decreased abundance and detected only in the control condition were grouped into six. Among the increased proteins, two clusters were involved in translation, mainly ribosomal proteins, as previously observed by GO analyses. Other clusters were related to the ribonucleoprotein complex, nuclear pore organization, and RNA inactivation/degradation **(Fig. 3)**.

**Fig. 3.**
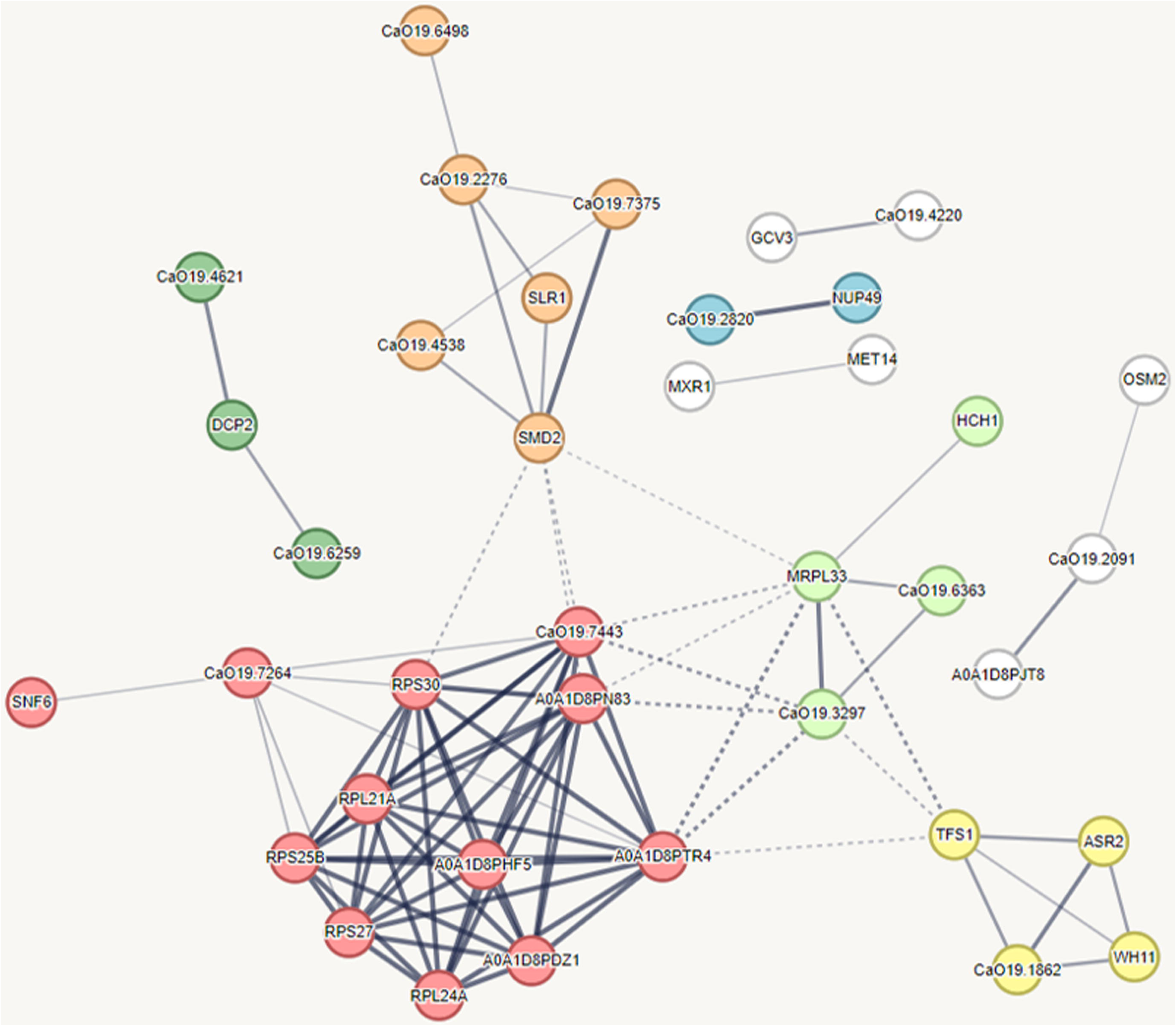
Networks of proteins with increased abundance in response to metformin. The line thickness between nodes (proteins) indicates the strength of data support, and the dotted lines indicate edges between clusters according to the STRING software. Clusters: translation (red and light green), ribonucleoprotein complex and RNA binding (orange), RNA inactivation/degradation (dark green), nuclear pore organization (blue), serine-type peptidase activity and heat shock protein (yellow), no significant enrichment detected (white).

Regarding proteins with decreased abundance, one of the main clusters was related to ATP synthesis-coupled electron transport. This cluster includes the proteins Mir1 (mitochondrial phosphate transporter), Cox5 (complex IV), and Cyt1 (complex III), as well as components of complex II (orf19.1480) and complex I (orf19.7590, orf19.4758) of the mitochondrial electron transport chain. There are two other remarkable clusters involved in ribosome biogenesis (small-subunit processome) and aminoacyl t-RNA biosynthesis. Clusters with chaperone activity and nucleocytoplasmic transport were also obtained **(Fig. 4)**.

**Fig. 4.**
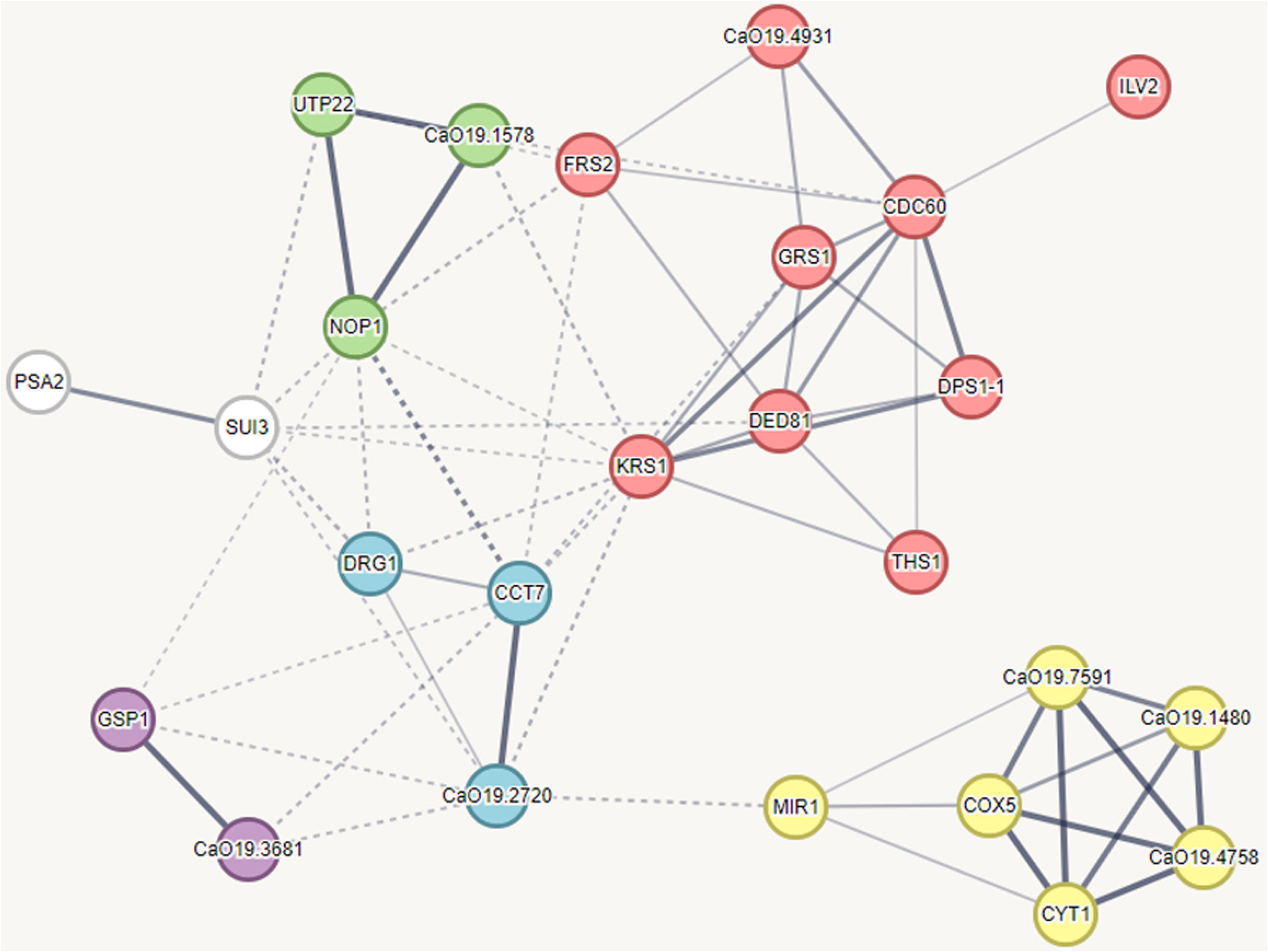
Networks of proteins with decreased abundance in response to metformin. The line thickness between nodes (proteins) indicates the strength of data support, and the dotted lines indicate edges between clusters according to the STRING software. Clusters: ATP synthesis-coupled electron transport (yellow), aminoacyl t-RNA biosynthesis (red), ribosome biogenesis (green), chaperone activity (blue), and nucleocytoplasmic transport (purple); no significant enrichment was detected (white).

### Analysis of the main processes affected by metformin

#### Inhibition of translation

Gene Ontology analysis and protein network studies have shown that a high number of proteins involved in translation are affected by metformin **(Supplementary Table S7)**. Of these, 28 increased and 16 decreased in abundance after treatment. Almost half of the proteins with increased abundance were ribosomal proteins, and more than half of those with decreased abundance were aminoacyl-tRNA synthetases, 75% of which were encoded by essential genes, in concordance with the decrease in *C. albicans* viability after treatment.

To study the inhibition of the translation process, we treated *C. albicans* cells with puromycin, a translation inhibitor in both prokaryotes and eukaryotes. As shown in **Fig. 5**, *C. albicans* SC5314 growth is inhibited in a concentration-dependent manner by puromycin in complete RPMI medium and strongly inhibited (approximately 75% of inhibition) with 0.125 mM puromycin. When metformin was added to compete with this concentration of puromycin, the OD increased with increasing concentrations of metformin to an intermediate level between the effects of puromycin and metformin alone, and in all cases, similar to that obtained with 50 mM of metformin. Thus, the puromycin effect is partially counteracted by metformin at concentrations as low as 6.25 mM, supporting the inhibition of translation as part of the anti-*Candida* effect of metformin.

**Fig. 5.**
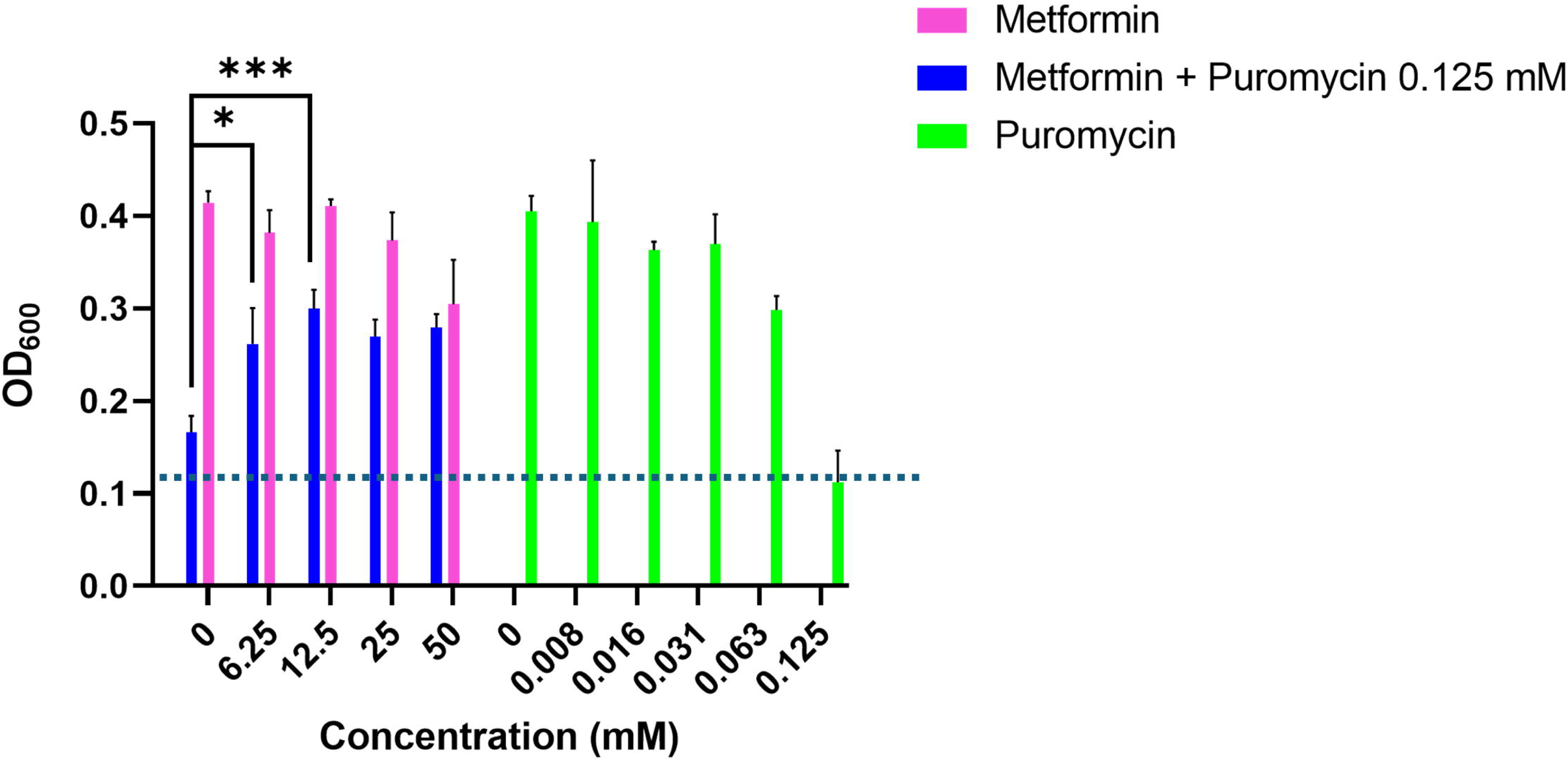
Sensitivity to puromycin and metformin in complete RPMI culture medium in microdilution plates. Increasing concentrations of puromycin 0-0.125 mM (green), metformin 0-50 mM (pink), and metformin + 0.125 mM puromycin (blue). The dotted line indicates the effect of puromycin 0.125 mM on *C. albicans* SC5314 growth, measured by OD at 600 nm. Data shown are the mean of three experimental replicates, with error bars representing standard deviation; \**p* < 0.05, \*\*\**p* < 0.001, unpaired t-test.

### Decrease in intracellular ATP levels

Proteins related to the GO Term ATP synthesis coupled electron transport formed one of the main clusters of proteins with decreased abundance after metformin treatment. Thus, intracellular ATP levels after 50 mM metformin treatment were measured, showing a significant decrease **(Fig. 6)**, indicating an alteration in mitochondrial function.

**Fig. 6.**
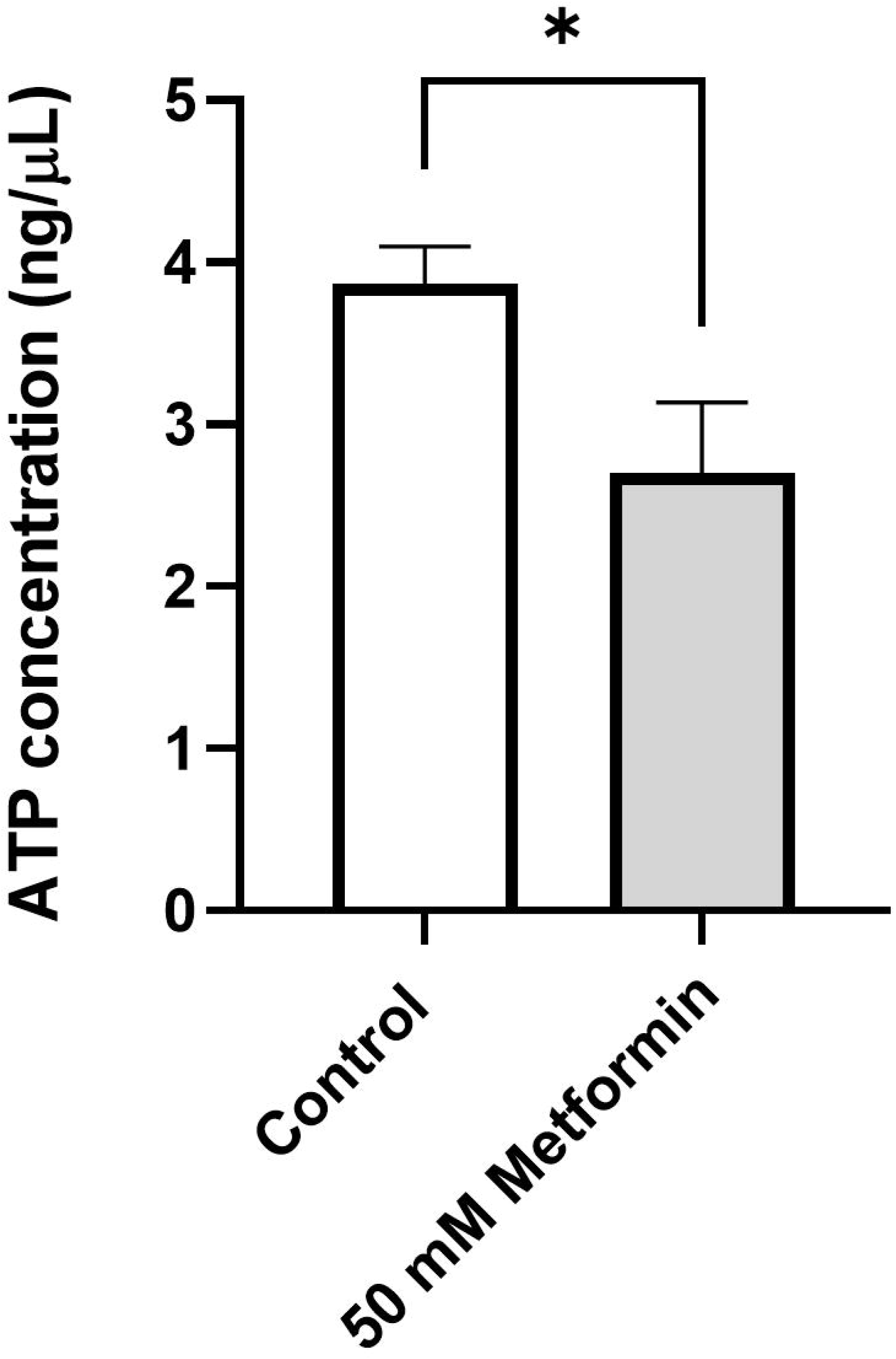
Effect of metformin on intracellular ATP levels. Intracellular ATP concentrations in cells growing with 50 mM metformin and untreated cells (control). Data shown are the mean of three experimental replicates, with error bars representing standard deviation; \**p* < 0.05, unpaired t-test.

### Effect on routes related to C. albicans filamentation, osmotic and oxidative stresses

The effect of metformin was markedly higher in RPMI complete medium under filament-inducing conditions. In addition, proteins related to dimorphic transition also significantly changed their abundance in the presence of metformin, most of which were involved in the increase in filamentous growth **(Supplementary Table S8)**. Thus, we decided to study the effect of metformin on several hyper-filamentous mutants. Mutants were chosen based on the studies conducted by [44] with the *C. albicans* homozygous deletion library constructed by Noble et al. (2010). Thus, mutants with the highest filamentous growth in the YPD liquid medium without induction were selected **(Fig. 7A)** and the effect of increasing concentrations of metformin on their growth was studied in complete RPMI medium. As observed in **Fig. 7B**, the mutants *hog1*Δ, *pbs2*Δ, and *ssk2*Δ, corresponding to the mitogen-activated protein kinase (MAPK) of the high osmolarity glycerol (HOG) signaling pathway, were more sensitive to metformin than the SC5314 strain. In contrast, the *cpp1*Δ mutant, a MAPK phosphatase that negatively regulates the Cek1 signaling pathway, was as sensitive as the wild-type strain. Increased sensitivity with respect to the wild type was not observed in other hyper-filamentous mutants not related to the HOG pathway or in mutants of Cek1 and Mkc1 MAPKs **(Supplementary Fig. S2)**. Thus, the HOG MAP-kinase pathway seems to be important for the survival of *C. albicans* in the presence of metformin. This pathway is involved in the response to oxidative stress. Some proteins involved in oxidative stress response were also detected in our proteomic study: Hat2 (histone acetyltransferase), Pst2 (oxidoreductase; fungal-specific), Hsp21 (heat shock proteins), and Mxr1 (oxidoreductase), with increased abundance after treatment with 50 mM metformin; Grx3 (glutaredoxin) and Ndt80 (transcription factor) that were only detected after exposure to metformin; and Gpx2 (oxidoreductase), orf19.4503 (DNA binding protein), orf19.7590 (oxidoreductase; complex I of the mitochondrial electron transport chain) with decreased abundance after treatment.

**Fig. 7.**
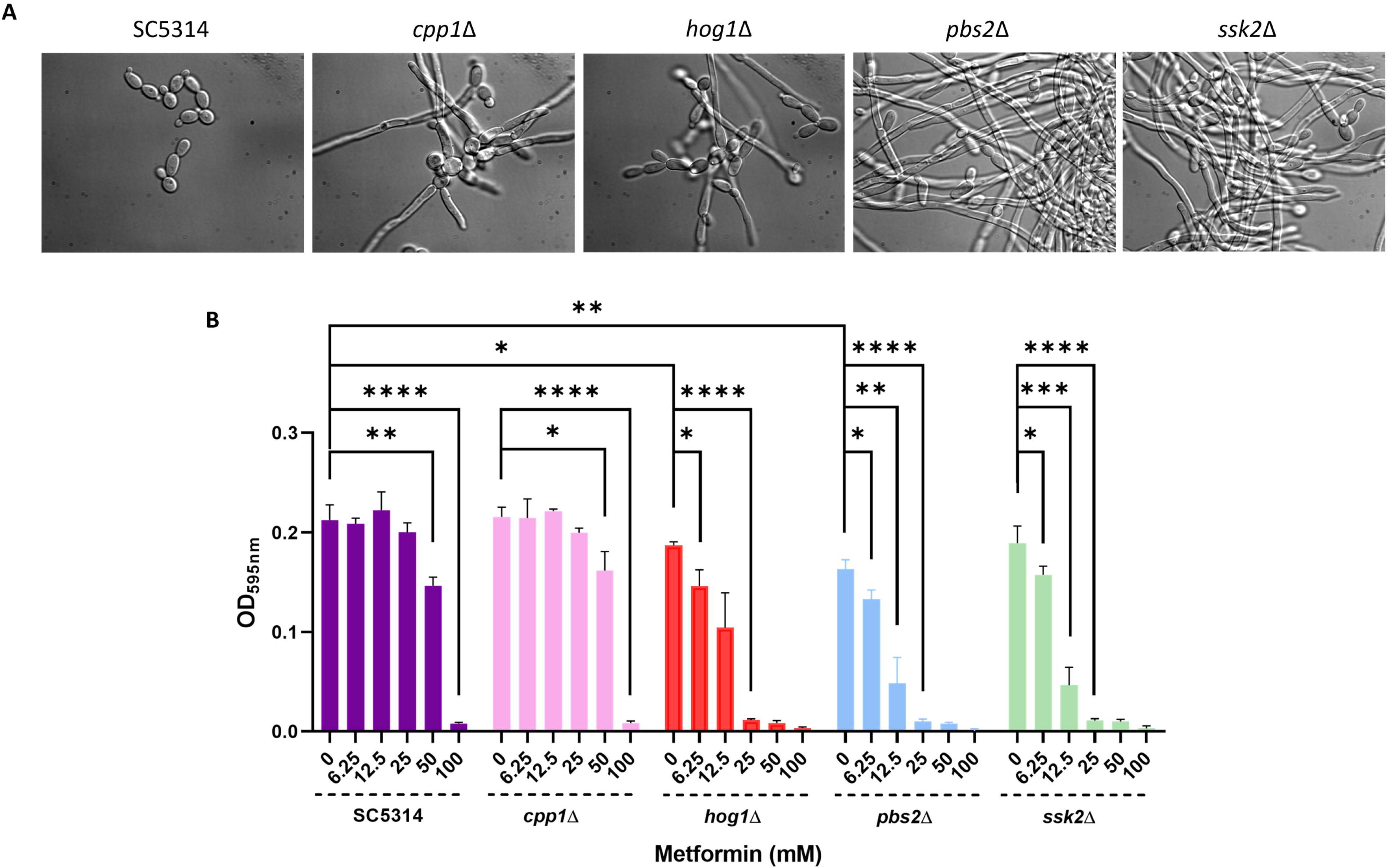
Effect of metformin on hyper-filamentous mutant strains. **(A)** Cell morphology of hyper-filamentous mutants observed by microscopy after growing the cultures at 37°C for 2 h in YPD medium. **(B)** Sensitivity of hyper-filamentous mutants to metformin in microdilution plates after treatment with increasing concentrations of metformin in complete RPMI medium; \**p* < 0.05, \*\**p* < 0.01, \*\*\**p* < 0.001, \*\*\*\**p* < 0.0001, unpaired t-test.

Consequently, the effects of the antioxidant agents N-acetylcysteine (NAC) and glutathione (GSH) were studied under our treatment conditions. As shown in **Fig. 8A**, NAC and GSH neutralized the effect of 100 mM metformin on the viability of the cells.

**Fig. 8.**
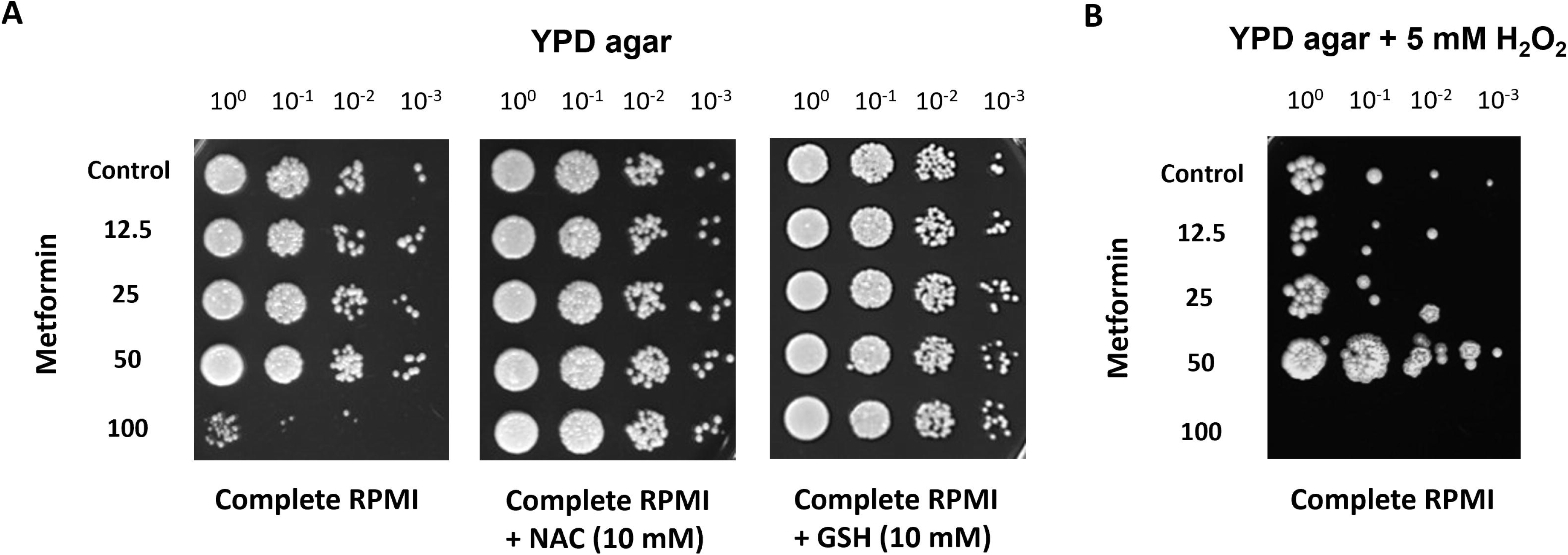
Effect of antioxidant agents on metformin activity. **(A)** Cell viability of ten-fold serial dilutions on YPD agar after treatment with increasing concentrations of metformin in complete RPMI, metformin + 10 mM NAC, or metformin + 10 mM GSH in microdilution plates. Effect of metformin pretreatment on viability in the presence of 5 mM H_2_O_2_ **(B)** Cell viability of ten-fold serial dilutions on YPD agar supplemented with 5 mM H_2_O_2_ after treatment with increasing concentrations of metformin in complete RPMI, in microdilution plates.

In addition, as the HOG MAPK pathway is also involved in the osmotic stress response, the effect of increasing concentrations of metformin pretreatment on resistance to different oxidative and osmotic stresses was analyzed. Surprisingly, only 50 mM metformin pretreatment increased the resistance of *C. albicans* to 5 mM hydrogen peroxide **(Fig. 8B)**. The protective effect was not observed with menadione and was very soft with osmotic agents **(Supplementary Fig. S3)**.

### Link with the response to antifungal drugs

Many proteins related to the response to antifungal drugs significantly changed in abundance after metformin treatment. The proteins described as changed in response to azoles (according to the data in the *Candida* Genome Database) were compared to the changes in response to metformin to correlate them with the possible intensifying effect. As shown in **TABLE 5**, the changes induced by metformin are possibly strengthening the azole effect. Possible interactions with other antifungal drugs are presented in **Supplementary Table S9**.

**TABLE 5.** Similarities and differences in gene expression and protein abundance in response to azole and metformin treatment.

| Antifungal | Antifungal response | Protein | Description | Metformin/Control Ratio log<sub>2</sub> |
| --- | --- | --- | --- | --- |
| Azoles | R ↓ | Pyc2 | Putative pyruvate carboxylase | 1.47 |
|  | ↑ | Hch1 | Ortholog of *S. cerevisiae* Hch1, a regulator of heat shock protein Hsp90 | 1.54 |
|  | ↑ | Ino1 | Inositol-1-phosphate synthase | 3.51 |
|  | ↑ | Wh11 | White-phase yeast transcript | 3.70 |
|  | R ↑ | Asm3 | Putative secreted acid sphingomyelin phosphodiesterase | -4.55 |
|  | R ↑ | Mir1* | Putative mitochondrial phosphate transporter | -1.73 |

\*Proteins encoded by essential genes. R, azole-resistant isolates. ↑, gene expression induced by azoles. ↓, gene expression repressed by azoles.

Sensitivity to the antifungal agent fluconazole was tested in combination with 50 mM metformin in complete RPMI medium (0.2% glucose) **(Fig. 9A)**, confirming previous results [10], which showed that metformin increases the antifungal effect of fluconazole. Additionally, the enhancement can be observed in the agar diffusion assay, where the confluence between fluconazole and metformin enlarges the fluconazole inhibition halo **(Fig. 9B)**.

**Fig. 9.**
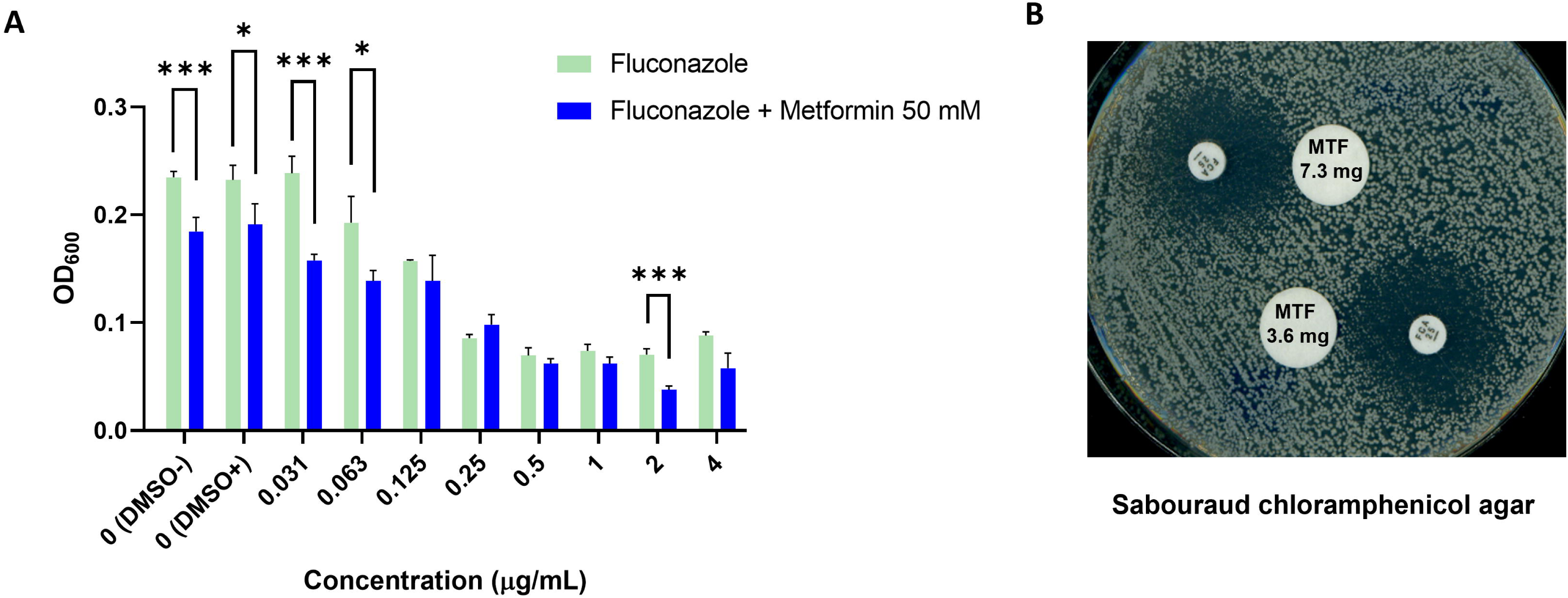
Effect of metformin on fluconazole activity. **(A)** Sensitivity to increasing concentrations of fluconazole 0-4 µg/mL (green) and fluconazole + metformin 50 mM (blue) in complete RPMI culture medium in microdilution plates. The amount of DMSO added as a control was the highest amount used with fluconazole. Data shown are the mean of three experimental replicates, with error bars representing standard deviation; *p < 0.05, ***p < 0.001, unpaired t-test. **(B)** Sabouraud chloramphenicol agar diffusion assay of fluconazole 25 µg (small disk) and metformin (MTF) at 7.3 mg and 3.6 mg.

## Discussion

In the context of drug repurposing, the antihyperglycemic drug metformin has emerged as a potential agent against *Candida* species [10, 11]. However, the effects of these compounds on *C. albicans* have not yet been revealed. Thus, this study delves into this matter using proteomics.

Metformin has shown an important impact on *C. albicans* growth and survival. First, we observed growth inhibition at a concentration of 50 mM. It also decreased filamentation, agar invasion abilities, and biofilm matrix at concentrations higher than 12.5 mM **(Fig. 1)**. The differences between our results and those previously reported [10] may be due to the freshly prepared metformin solutions used for the present work, the culture media, and the glucose concentration used. Our experiments have shown more efficient inhibition of growth under conditions that induce *C. albicans* filamentation (complete RPMI *vs.* YPD) and low glucose concentration (0.2% *vs.* 2%) **(Supplementary Fig. S1)**. The MIC_50_ calculated for metformin against *C. albicans* by Xu et al. (2018) in RPMI-MOPS medium with 2% glucose was 120 mM (20.41 ± 0.32 mg/mL), more than twice our 50 mM. The reduction in the effect of metformin when high amounts of fermentable sugars are available in the medium has also been observed in *Saccharomyces cerevisiae* growth [41], although *C. albicans* is *Crabtree-negative*. The influence of glucose concentration is extensible to other metformin effects, such as the increase in lifespan of *Caenorhabditis elegans* and the antibacterial action on *Escherichia coli* [40]. This effect might be partially explained by the activation of AMPK, which has been described as one of the main targets of metformin under glucose starvation conditions [20]. In *C. albicans*, the AMPK homologue is Snf1, which is phosphorylated by Sak1 in response to nutrient stress, such as low glucose levels [45, 46]. This phosphorylation has been shown critical for *C. albicans* use of alternative carbon sources, maintenance of cell integrity, invasive growth, and regulation of the yeast-to-hypha transition. In fact, mutants in *SNF1* or *SAK1* have a similar phenotype to metformin-treated cells with respect to filamentous and invasive growth [46]. To show the involvement of AMPK activation in the metformin effect on *C. glabrata,* Xu et al. (2018) treated this yeast with the AMPK agonist AICAR (5-Aminoimidazole-4-Carboxamide-1-Beta-D-Ribofuranoside) and demonstrated a small but statistically significant reduction in metabolic activity, suggesting that AMPK-dependent mTOR inhibition can contribute to growth reduction. In the same study, rapamycin (mTOR inhibitor) showed some growth inhibition, but without synergism with metformin, suggesting that alternative metabolic pathways to mTOR may also be involved in biguanide activity. In addition, the mitochondrial complex I inhibitor rotenone did not inhibit the growth of *C. glabrata*. In our case, *C. albicans* SC5314 Snf1 was not phosphorylated at several time points (up to 1h) in RPMI with 50 and 100 mM metformin (data not shown). Therefore, there is more to discover regarding the anti-*Candida* action of metformin.

Based on our results, we chose RPMI (0.2% glucose) and a sublethal concentration of 50 mM metformin for proteomics and other studies. This concentration exceeds those achieved in blood under regular metformin treatment and those used with human cell lines [47], but it is also the range of effective concentrations observed by other authors [41]. Metformin concentrations in the jejunum peak at 500 µg/g, up to 300 times higher than in plasma, as mentioned before [48], and can cause changes in the microbial composition [49], which might include intestinal *Candida* cells. It has been postulated that in susceptible patients (immunodepressed, with solid cancer, and recovering from abdominal surgery), the source of invasive candidiasis could be the gastrointestinal tract [4-6]. A recent study showed that patients with allogenic hematopoietic cell transplants who suffered from fungal bloodstream infections by *Candida* experienced a prior marked intestinal expansion of *Candida* species [50]. Therefore, the anti-*Candida* effects of metformin on viability and virulence factors, such as filamentation and biofilm formation, could be clinically interesting [51]. In addition, the potentiation of other widely used antifungals could help in the treatment of these infections.

The quantitative proteomic study allowed us to identify and quantify 1899 proteins. Of them, 95 showed increased abundance, while 47 decreased after metformin treatment. Many of the proteins that were significantly more abundant after treatment, and some of the less abundant ones, were proteins with unknown functions. It is also worth highlighting that 26 of the proteins with decreased abundance were encoded by an essential gene according to the data in the *Candida* Genome Database and Segal et al. (2018), a sign of the intense effect on *C. albicans* physiology, and probably responsible for the *C. albicans* lethality at high concentrations.

The functions of the proteins were studied using the GO Term Finder tool from the *Candida* Genome Database. GO enrichment in the biological process “translation” included the largest group of proteins altered by metformin treatment **(Supplementary Table S6)**. A reduction in protein synthesis has also been observed in multiple myeloma cells treated with the antihypertensive drug syrosingopine and metformin, but not with single agents [52]. In a recent study on the effect of this drug on JHH-7 cells, the authors observed marked suppression of global protein ubiquitination and concurrent inhibition of both protein synthesis and degradation with 10 mM metformin [25]. In addition, the inhibition of protein synthesis is common to other stress responses [53].

In our study, most of the proteins with significantly decreased abundance were aminoacyl-tRNA synthetases encoded by essential genes, whereas most of those with significantly increased abundance were ribosomal proteins **(Fig. 3-4)**. Aminoacyl-tRNA synthetases play a central role in protein synthesis and have garnered interest as potential antimicrobial targets [54]. Ribosomal proteins (RPs) are essential for ribosome assembly and function [55]. It has been proposed that the accumulation of RPs can arise from defects in ribosome assembly caused by an imbalance between RPs and rRNA [56, 57]. Puromycin is an antibiotic that is structurally similar to aminoacylated tRNA. It produces a premature termination of translation when incorporated into nascent polypeptide chains, leading to ribosomal subunit dissociation [58], effects very similar to those induced by metformin. Thus, we examined its effects on *C. albicans* SC5314 and observed that 0.125 mM puromycin strongly inhibited *C. albicans* growth. When different concentrations of metformin were added, the puromycin effect was neutralized **(Fig. 5)**. Thus, both drugs interfere, showing the inhibition of translation as part of the anti-*Candida* effect of metformin.

In addition, the decrease in ATP levels **(Fig. 6)** affects protein synthesis and turnover [25]. Other proteins affecting translation are worth mentioning, such as Mss51, a putative mRNA maturation factor [59, 60], which increases in the presence of metformin and is fungal-specific (no human or murine homolog); thus, it might be interesting as a fungal target. Another interesting protein can be Rpn11, involved in proteasome and mitochondrial fission [61, 62].

The abundance of proteins related to antifungal drug response also significantly changed. It is important to underline that the proteins whose transcripts have been described as overexpressed in azole-resistant isolates suffer a reduction by metformin treatment **(Table 5)**. These reductions might explain the reinforcement of azole treatment by metformin **(Fig. 9** and [10]). This is the case for Asm3 [63], the second protein with the highest decrease in relative abundance upon metformin treatment **(Table 3)**, which is described as a sphingomyelin phosphodiesterase involved in ceramide formation. This is interesting because fungal sphingolipids have emerged as potential targets for novel antifungal agents [64]. Or the protein Mir1 [65], encoded by an essential gene that transports inorganic phosphate into the inner matrix of fungal mitochondria, where it is used by ATP synthase to generate ATP, and has also been proposed as an antifungal target [66]. Metformin also increases the abundance of proteins whose transcripts are repressed in an azole-resistant isolate, such as Pyc2 [63], described as a pyruvate carboxylase that catalyzes the ATP-dependent carboxylation of pyruvate to oxaloacetate [67].

Another protein showing decreased abundance upon metformin treatment is Erb1, which is also encoded by an essential gene with a predicted role in ribosomal large subunit biogenesis. The heterozygous *erb1* mutant is hypersensitive to flucytosine [68]. Thus, combined treatment with metformin may increase sensitivity to this antifungal agent. However, as Zhao et al. (2025) described antagonism between these two substances, more experiments are necessary to prove this point. Grx3, which increased in the presence of metformin, was also induced by flucytosine treatment and was related to cell stress [69]. It is a monothiol glutaredoxin (oxidoreductase) required for growth under low iron conditions [70], and this is in agreement with a previous observation in *S. cerevisiae*, in which the effect of metformin resembles iron deficiency [41]. We found other proteins (Gpx2, Cyt1, and Cox5) that decreased in abundance, consistent with this iron deficiency-like state [71, 72]. However, the addition of iron to the culture medium did not make *C. albicans* less sensitive to metformin (data not shown). Gpx2 is a glutathione peroxidase (oxidoreductase) involved in the oxidative stress response, and Cyt1 and Cox5 (oxidoreductase, encoded by an essential gene) are components of complexes III and IV of the mitochondrial electron transport chain. As previously published [19, 23], metformin inhibits complex IV, which is consistent with our results.

It has also been suggested [10] that metformin may inhibit efflux pumps and potentiate the effect of antifungals. In contrast, we observed an increase in the abundance ofNdt80, a *CDR1* activator, which was not detected under control conditions.

With respect to the dimorphic transition, 28 of the significantly changed proteins were involved in this process **(Supplementary Table S8)**, in concordance with the phenotypic observations **(Fig. 1)**. Thus, we studied the effect of metformin on the growth of mutants with defects in the dimorphic transition. Analyses of hyper-filamentous mutants revealed the importance of the Hog1 MAP-Kinase pathway for the survival of *C. albicans* in the presence of metformin **(Fig. 7)**. This pathway is involved in the responses to both oxidative and osmotic stress [73, 74], and *Candida*’s ability to resist oxidative stress is considered part of its resistance mechanisms to phagocytes, which constitute one of the main defenses against invasive candidiasis [75]. In contrast, the *cpp1D* mutant, which is also hyper-filamentous but not involved in the Hog1 pathway, showed the same level of sensitivity as the wild-type strain. Another protein related to Hog1 is Osm2, a putative mitochondrial fumarate reductase, and its mutant has a reduced ability to invade agar [76]. This protein increases twofold in the presence of metformin and also increases in the presence of nitrosative stress, cell wall damage, serum, low H_2_O_2_ concentrations [77], and DNA damage [78]. As Hog1 downregulates Osm2, the mutant strain should have this protein derepressed, as in the presence of metformin, and could increase the sensitivity to the drug in strains with defects in the Hog1 pathway. This finding highlights the importance of the Hog1 MAPK pathway in the response to metformin.

Ess1 is another interesting protein that increases in the presence of metformin. This protein is involved in the Cph1 pathway. The heterozygous mutant (coded by an essential gene) was not able to form filaments. This increase may be due to a compensatory mechanism for the decrease in filamentation.

The action of metformin also affects the oxidative stress response, as shown by the neutralization of *C. albicans* lethality produced by 100 mM metformin by the antioxidant agents N-acetylcysteine (NAC) and glutathione (GSH) **(Fig. 8A)**. In support of this, it has been shown that pretreatment with a low-level ROS-inducer can protect against further oxidative stress in *C. albicans* [79] and that is what we have observed when treating *C. albicans* with a sublethal concentration of metformin (50 mM) **(Fig. 8B)**. As the proteomic study has shown, several proteins involved in the oxidative stress response significantly changed their abundance after treatment with 50 mM metformin, mainly oxidoreductase proteins, such as the flavodoxin-like protein Pst2, which is induced by the main regulator of the oxidative stress response, Cap1, and heat shock proteins such as Hsp21, which is involved in the oxidative stress response [80]. This effect has not been observed with lower metformin concentrations, probably because the anti-ROS response was not as intensely induced.

Mitochondria are one of the main targets of metformin [19, 22], and several proteins with decreased abundance after metformin treatment were clustered in the group related to ATP synthesis-coupled electron transport **(Fig. 4)**. This is the case for the aforementioned protein Mir1, as well as proteins of complex I (orf19.7590 and orf19.4758) and complex IV (Cox5), previously mentioned, and other proteins from the other complexes. As published, inactivation of the mitochondrial protein Goa1, putative subunits of respiratory complex I, or factors required for mitochondrial biogenesis and morphology in *C. albicans*, all result in attenuated virulence in a mouse model of systemic candidiasis [81]. In addition, the abundance of proteins related to ATPase/ATP synthase activity significantly changed after the treatment. This is the case for orf19.7345 and orf19.5201.1, which are among the proteins with the highest increase in their relative abundance **(TABLE 1)**, orf19.3686, among the proteins exclusively detected in the treated yeasts **(TABLE 2)**, and Ena21, which was only detected in non-treated control conditions **(TABLE 4)**. As previously mentioned, a decrease in intracellular ATP levels was observed after the treatment with 50 mM metformin **(Fig. 6)** which points to an inhibition of mitochondrial functions caused by metformin.

Another interesting point has been described in a recent study: the involvement of metformin in autophagy [11]. These authors showed that metformin increases autophagy in planktonic cells while inhibiting it in biofilms. In our conditions (filamentation) and in the presence of metformin, Dcp2, an mRNA decapping factor that plays a crucial regulatory role in autophagy, assisting through autophagy the survival of *C. albicans* to DNA damage stress [82], increases 4x after the treatment, probably trying to help *C. albicans* survival to the stress of the treatment.

In conclusion, high concentrations of metformin, which might be achieved in the GI tract, seriously affect *C. albicans* viability and dimorphic transition. The proteomic study revealed a high number of proteins encoded by essential genes, the abundance of which decreased after treatment. Processes such as translation, mitochondrial function, ATP biosynthesis, and oxidative stress response are affected by metformin, as deduced from proteomic and functional studies. The pleotropic effects of metformin might explain the potentiation of the antifungal action of azoles or Amphotericin B. Thus, as metformin is one of the most prescribed drugs worldwide, its antifungal action should be studied in more depth.

## Data availability

Supplemental data. This article contains supplemental data.

## Author Contributions

Conceptualization, G.M. and C.G.; methodology, V.M., C.R., C.N., M.L.H., software, V.M. and M.L.H.; validation, V.M., C.R., M.L.H.; formal analysis, V.M., G.M. and M.L.H.; investigation, V.M., C.R., C.N., G.M. and C.G.; resources, G.M., and C.G.; data curation, V.M. and M.L.H.; writing—original draft preparation, V.M. and G.M.; writing—review and editing, V. M., G.M. and C.G.; visualization, V.M. and C.R.; supervision, G.M. and C.G.; project administration, G.M. and C.G.; funding acquisition, G.M., and C.G. All authors have read and agreed to the published version of the manuscript.

## Use of AI

The authors declare the use of AI tools only for basic checks of grammar, spelling and punctuation.

## Abbreviations

AMPK: AMP-activated protein kinase
GI: gastrointestinal tract
NAC: N-acetylcysteine
GSH: glutathione.

---

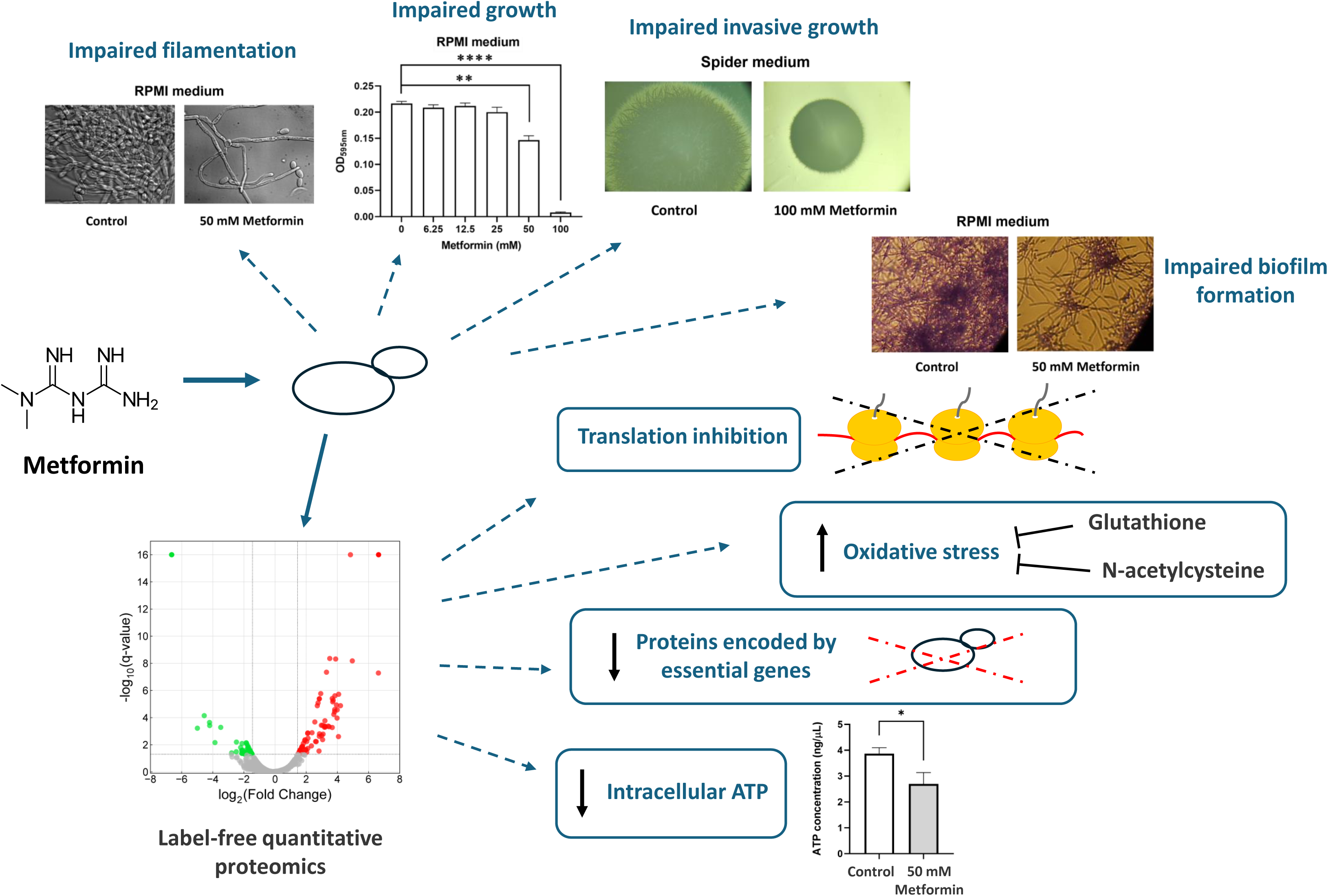

