## Supplementary figures S1-S3 for "Proteomics to uncover new actors in the antifungal action of metformin. Impact on virulence traits, oxidative stress, and essential proteins from *Candida albicans*"

### SUPPLEMENTAL MATERIAL

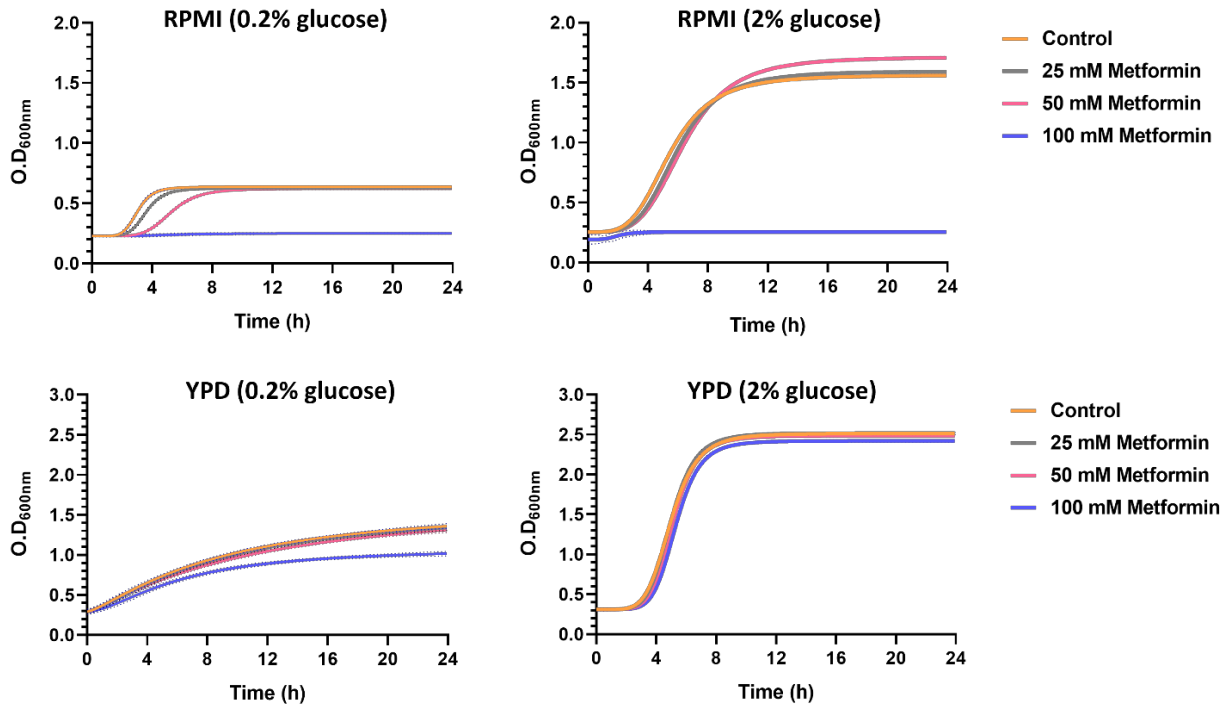

**Supplemental Figure S1. Glucose concentration influence on the inhibitory effect of metformin.** *C. albicans* growth curves in complete RPMI and YPD culture media with low (0.2%) and high (2%) amount of glucose and increasing concentrations of metformin. Data shown are the mean of three experimental replicates with error bars representing standard deviation.

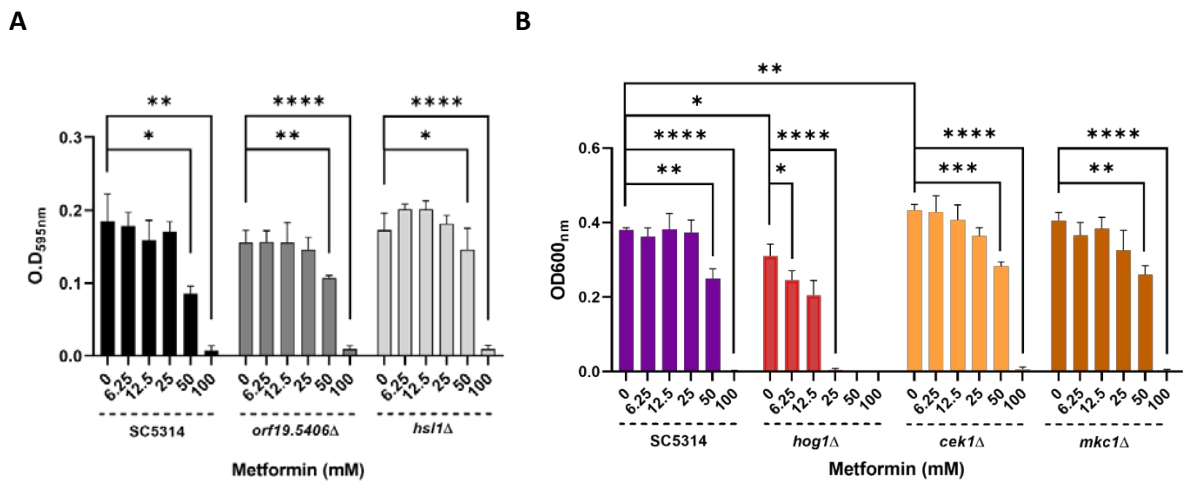

**Supplemental Figure S2. Effect of metformin on several mutants.** (A) Sensitivity to metformin of hyper-filamentous mutants not related to the HOG pathway in microdilution plates after a treatment with increasing concentrations of metformin in complete RPMI medium. (B) Sensitivity to metformin of MAPKs mutants in microdilution plates after a treatment with increasing concentrations of metformin in complete RPMI medium; \*$p < 0.05$, \*\*$p < 0.01$, \*\*\*$p < 0.0001$, unpaired t-test.

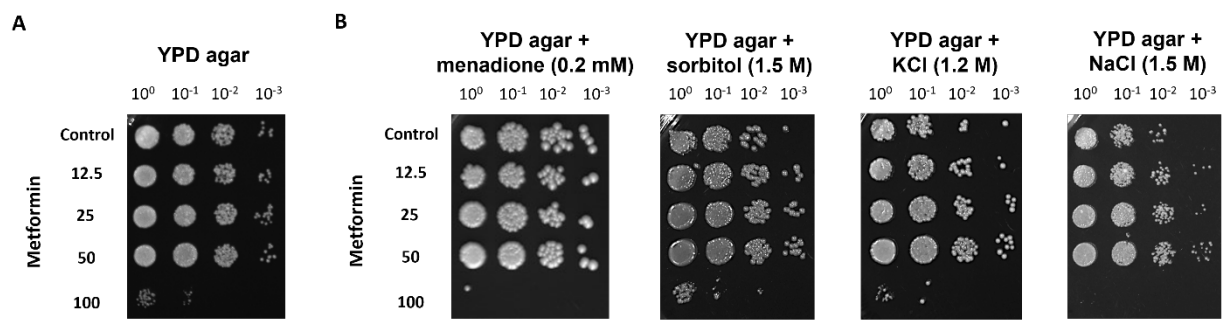

**Supplemental Figure S3. Effect of menadione and osmotic agents on metformin activity. (A)** Cell viability of ten-fold serial dilutions on YPD agar after a treatment with increasing concentrations of metformin in complete RPMI, in microdilution plates. **(B)** Cell viability of ten-fold serial dilutions on YPD supplemented with 0.2 mM menadione, 1.5 M sorbitol, 1.2 M KCl and 1.5 M NaCl after a treatment with increasing concentrations of metformin in complete RPMI, in microdilution plates.
